# Endogenous glucocorticoids are required for normal macrophage activation and gastric *Helicobacter pylori* immunity

**DOI:** 10.1101/2024.01.14.575574

**Authors:** Stuti Khadka, Sebastian A. Dziadowicz, Xiaojiang Xu, Lei Wang, Gangqing Hu, Jonathan T. Busada

## Abstract

Glucocorticoids are steroid hormones well-known for their potent anti-inflammatory effects. However, their immunomodulatory properties are multifaceted. Increasing evidence suggests that glucocorticoid signaling promotes effective immunity and that disruption of glucocorticoid signaling impairs immune function. In this study, we conditionally deleted the glucocorticoid receptor (GR) in the myeloid lineage using the *LysM-Cre* driver (myGRKO). We examined the impact on macrophage activation and gastric immune responses to *Helicobacter pylori*, the best-known risk factor of gastric cancer. Our results indicate that compared to WT, GRKO macrophages exhibited higher expression of proinflammatory genes in steroid-free conditions. However, when challenged in vivo, GRKO macrophages exhibited aberrant chromatin landscapes and impaired proinflammatory gene expression profiles. Moreover, gastric colonization with *Helicobacter* revealed impaired gastric immune responses and reduced T cell recruitment in myGRKO mice. As a result, myGRKO mice were protected from atrophic gastritis and pyloric metaplasia development. These results demonstrate a dual role for glucocorticoid signaling in preparing macrophages to respond to bacterial infection but limiting their pathogenic activation. In addition, our results support that macrophages are critical for gastric anti- *Helicobacter* immunity.

**Graphical Abstract:** 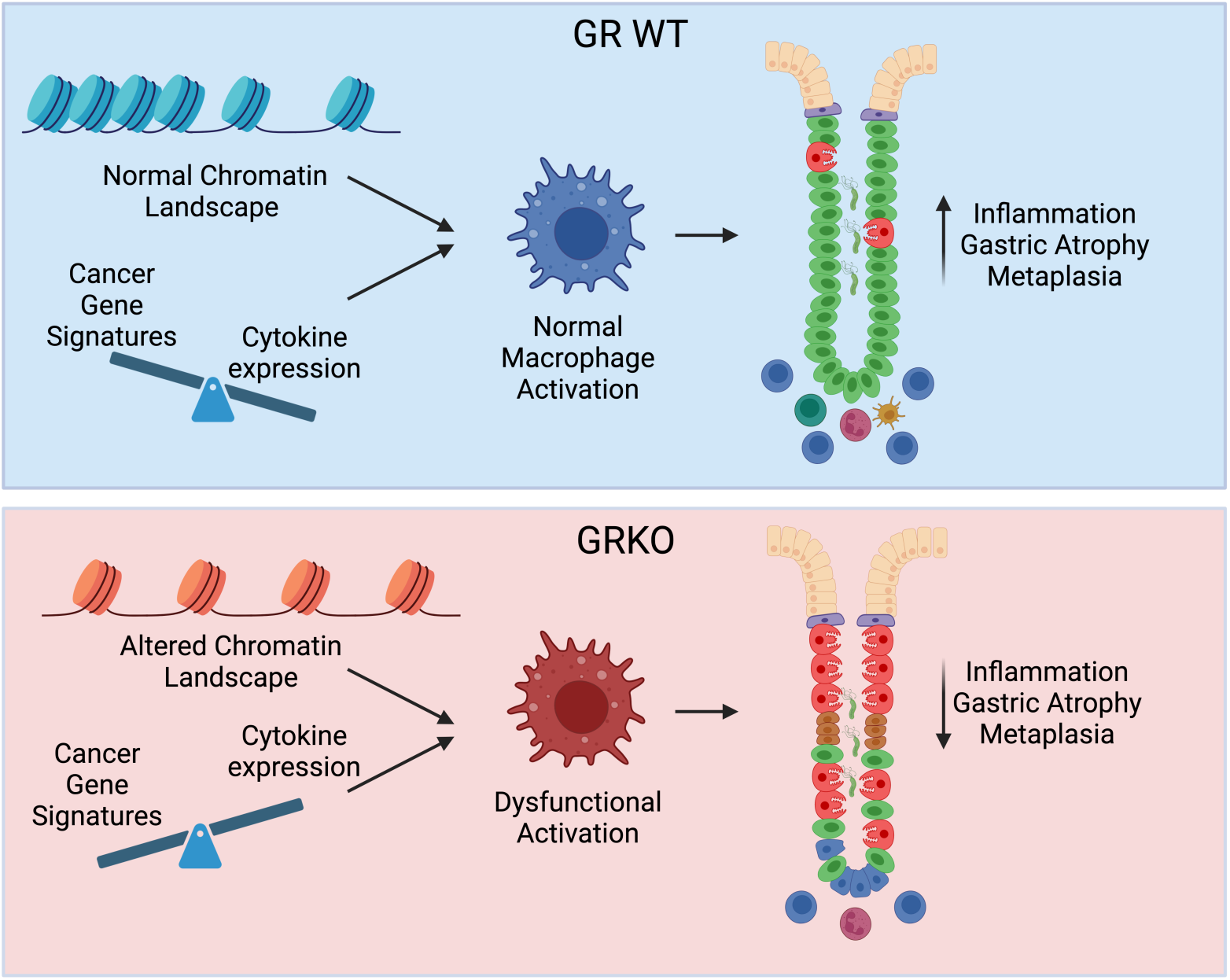

## Introduction

Glucocorticoids are steroid hormones produced as the end product of the hypothalamic-pituitary-adrenal axis. They signal through the glucocorticoid receptor (NR3C1, hereafter GR), a ligand-dependent transcription factor, regulating a wide array of cellular and physiological functions. Clinically, glucocorticoids are widely used to treat a host of inflammatory disorders where they limit the activation and cytokine production of pathogenically activated immune cells^(1)^. Their common clinical use has contributed to a dogmatic perception that glucocorticoids are purely anti-inflammatory. However, glucocorticoids exert diverse immunomodulatory effects, both inhibiting and enhancing aspects of immunity ^(2)^. Recent studies have found that signaling by glucocorticoids enhance T cell antigen specificity, regulate B cell homing, and enhance macrophage responses to lipopolysaccharide (LPS) ^(3–6)^. However, the role of glucocorticoids in supporting normal immune functions are poorly understood.

Gastric cancer is the 5th most common cancer worldwide ^(7)^. Chronic infection by the bacterium *Helicobacter pylori* is the highest known risk factor of gastric cancer. *H. pylori* infection is extremely prevalent, affecting approximately 50% of the global population. The infection causes chronic inflammation within the gastric mucosa, initiating a well-established histopathological cascade of gastric atrophy and metaplasia, eventually resulting in gastric adenocarcinoma ^(8)^. Gastric inflammation is ubiquitous among *H. pylori-infected* individuals, but only a small portion of infected individuals will develop gastric cancer ^(9)^. Inflammation is important for controlling gastric *H. pylori* burden, but more intense immune responses also drive gastric atrophy and may accelerate gastric cancer initiation ^(10)^.

Macrophages are critical for multiple facets of the immune response. Early responses from tissue-resident macrophages detect pathogen-associated molecular patterns (PAMPs) and release chemokines to attract other immune cells. Bacterial phagocytosis and cytokine release reduce bacterial burden but may also induce off-target damage to host cells. Moreover, macrophage antigen presentation is important for generating T cell responses. Numerous studies have demonstrated that macrophages are essential during the anti-*Helicobacter* immune response. Macrophages control *H. pylori* gastric burden, and impaired macrophage function leads to higher *H. pylori* bacterial loads ^(11)^. However, macrophages also promote gastric atrophy progression and macrophage depletion reduces *H. pylori*-associated gastric atrophy ^(12)^. Macrophage-derived cytokines also have been implicated in promoting pyloric metaplasia (PM) development (a putative preneoplastic lesion), and macrophage depletion prevents PM development following gastric injury ^(13, 14)^. Thus, enhance macrophage activation may promote gastric atrophy and confer increased gastric cancer risk.

We recently reported that glucocorticoids are master regulators of gastric inflammation and that systemic disruption of glucocorticoid production by adrenalectomy caused spontaneous gastric inflammation, leading to gastric atrophy and PM development ^(15)^. Macrophages were key drivers of these gastric pathologies, as macrophage depletion prevented gastric atrophy and metaplasia. However, GR expression is ubiquitous in the stomach, and the cell-intrinsic effects of glucocorticoid signaling on macrophages and their impact on gastric immunity are unclear. In this study, we utilized the Cre-Lox system to specifically delete the GR from the myeloid lineage (myGRKO) and investigated the subsequent impact on macrophage activation and anti-*H. pylori* immunity. We found that GRKO macrophages were dysfunctional, exhibiting aberrant chromatin accessibility and transcriptional responses following bacterial challenge. Moreover, myGRKO mice exhibited impaired gastric inflammation and reduced gastric atrophy following infection challenge with either *H. pylori* or *H. felis.* These results demonstrate a critical role for glucocorticoids in promoting normal proinflammatory macrophage function and that macrophages play a crucial role in coordinating gastric anti-*Helicobacter* immune responses.

## Material and Methods

### Animal care and Treatment

Mice were housed and maintained in a temperature and humidity-controlled vivarium with 12-hour light/dark cycles and provided with standard chow and water *ab libitum*. The GR was deleted from the myeloid lineage by crossing *Nr3c1^flox/flox^* (Jax Stock # 021021) and homozygous *LysM-Cre* (Jax Stock # 004781) were purchased from the Jackson Laboratories. *Nr3c1^flox/flox^*; *LysM^+/Cre^* mice were considered myeloid GRKO (myGRKO) while *Nr3c1^flox/flox^; LysM^+/+^* littermates were used as WT controls. Both sexes were used for these studies. Routine genotyping was performed by TransnetYX. For gastric colonization with *Helicobacter,* mice were inoculated with 500 μL of brucella broth containing 10^9^ CFU of either *H. pylori* or *H. felis* two times 24 hours apart. For peritoneal macrophage isolation, mice received a single intraperitoneal (IP) injection of 1mL sterile 3% Brewers Thioglycollate media (Sigma-Aldrich). After 4 days, peritoneal lavage was plated, and nonadherent cells were removed by washing with prewarmed 1xPBS.

### Bacterial preparation

The *H pylori* PMSS1 strain ^(16)^ (a gift from Manuel Amieva, Stanford University) and *H. felis* CS1 strain (ATCC 49179) were grown on tryptic soy agar plates (BD Biosciences) with 5% defibrinated sheep blood (Hemostat Labs) and 10 mg/mL vancomycin (Alfa Aesar) under microaerophilic conditions at 37°C for 2 days. Bacteria were harvested and transferred to Brucella broth containing 5% fetal bovine serum and 10 mg/mL vancomycin and grown overnight at 37°C under microaerophilic conditions with agitation. Bacteria were centrifuged and resuspended in fresh Brucella broth without antibiotics before spectrophotometry. Bacterial number were determined by a standard curve of bacterial counts and absorbance at OD 600. Gram staining was used to confirm culture purity.

### Tissue preparation

Mice were euthanized by cervical dislocation without anesthesia. Stomach tissue was processed for flow cytometry, immunostaining, and RNA isolation as previously described ^(17)^ (17). Briefly, stomachs were opened along the greater curvature and washed in phosphate buffered saline (PBS) to remove the gastric contents. One half of the gastric corpus was used for flow cytometry. A 4 mm biopsy punch of the gastric corpus greater curvature was collected for RNA isolation and immediately snap-frozen in liquid nitrogen. From the other half of the stomach, a biopsy was collected to confirm *Helicobacter* colonization and the remainder was fixed overnight in 4% paraformaldehyde at 4oC. After fixation, stomachs were washed with PBS and cut into strips. A strip was put in 30% sucrose, embedded in optimal cutting temperature compound for cryosectioning and another strip was transferred in 70% ethanol for routine processing and paraffin embedding.

### Immunofluorescence staining

Five µm thick cryosections from the gastric corpus greater curvatures were stained for key epithelial cells and metaplasia markers. The sections were incubated with H+/K+ ATPase (MBL International), MIST1 (Cell Signaling Technologies), or CD44v9 (Cosmo Bio) overnight at 4°C. Primary antibody was omitted as a negative control. Sections were incubated in secondary antibodies for 1 hour at room temperature. Fluorescent conjugated *Griffonia simplicifolia* lectin (GSII; ThermoFisher Scientific) was added with secondary antibodies. Sections were mounted with Vectastain mounting media containing DAPI (Vector Laboratories). Images were obtained using a Zeiss 710 confocal laser-scanning microscope (Carl-Zeiss GmbH) and running Zen Black imaging software.

### Flow Cytometry

Single cell suspensions were collected from the gastric corpus. Stomachs were washed in 1X PBS, minced, and incubated in Hanks Balanced Salt Solution containing 5% fetal bovine serum and 5 mM EDTA on a shaker incubator at 37°C. The tissue was then incubated with DNAse (0.5mg/ml, Worthington Biochemical) and Collagenase Type IV (1mg/ml, Worthington Biochemical) and disassociated by pushing through a 100 µM cell strainer. Debris was removed using a 40% OptiPrep gradient (Serumwerk). Cells were stained for 20 minutes at 4°C with CD45 (Clone 104), B220 (Clone RA3-6B2), CD3e (Clone 145-2C11), CD11b (Clone M1/70), F480 (Clone BM8), MHCII (Clone M5/114.15.2), Ly6G (1A8) (all from Bioledgend) and SiglecF (Clone E50-2440 eBiosciences). Actinomycin D (Invitrogen) was added to label dead cells. Samples were analyzed on a BD Fortessa (BD Bioscience). The data was analyzed using Cytobank (Beckman Coulter).

### ATACseq

Mice were IP injected with sterile Thioglycollate medium as described above. After four days, mice received a single IP injection of 10 million *H. pylori* in 500 ul of sterile saline. After 3 hours, peritoneal lavage was collected, and the cells were stained with CD45 (Cone 104), CD11b (Clone M1/70), F480 (Clone BM8), and CX3CR1 (SA011F11) for 20 minutes at 4oC. Actinomycin D (ThermoFisher) was used to label dead cells. Live CD45+CD11b+F480+CX3CR1+ macrophages were sorted on a FACS Aria III (BD Bioscience). ATACseq was performed using 50,000 cells. ATAC libraries were prepared using the Tagment DNA Enzyme and Buffer Kit (Illumina) following the Omni-ATAC protocol ^(18)^. Libraries were sequenced to 40 million reads on a Nextseq 2000 (Illumina) by the Marshall University Genomics and Bioinformatics Core facility. ATAC-seq data analysis was described previously^(19)^. Briefly, ATAC-seq reads were aligned to the mouse reference genome (mm10) by Bowtie 2 ^(20)^, ATAC-seq read enriched regions were identified by macs 3 ^(21)^, differentially accessible regions (DAR) by EdgeR 3 ^(22)^, target genes of DARs by GREAT (version 4.0.4, Stanford University) ^(23)^, and motif analysis for DARs by HOMER ^(24)^. Pathway analysis for target genes of DARs were conducted by Metascape ^(25)^. A *P-*value cutoff of 0.05 was used for pathway and motif analysis to assess the subtle changes between unstimulated WT and GRKO macrophages. A q-value cutoff of 0.01 was used for the *H. pylori-stimulated* groups.

### RNAseq

WT and GRKO peritoneal macrophages were stimulated and isolated as described above in the ATACseq section. RNA was isolated using the Microelute RNA isolation kit (Omega Bio-Tek) with on-column DNase treatment (Omega Bio-Tek) following the manufacturer’s protocol. Libraries were prepared by the West Virginia University Genomics Core using the NEBNext Low Input/Single Cell RNA-Seq kit (New England Biolabs). Libraries were sequenced to 30 million reads on a Nextseq 2000 (Illumina) by the Marshall University Genomics and Bioinformatics Core facility. RNA-Seq data analysis followed procedures as previously described ^(19)^. RNA-Seq read were aligned by subread v2.0.1 ^(26)^. Read summarization was performed by RefSeq and transcript annotation by Featurecounts ^(27)^, and differentially expressed genes identified by EdgeR3 ^(22)^. Gene Set Enrichment Analysis (GSEA 4.3.2) was used to identify pathway annotation of the genes preferentially expressed in a test group compared to the reference group. For each comparison, a “rnk” file was prepared with the expressed genes (RPKM≥3) and their corresponding log Fold Change and run against the KEGG and Reactome pathway collections pre-defined in MSigDB ^(28)^. The enrichment score on the GSEA was calculated using 1000 permutations. The resulting GSEA pathways were further analyzed and visualized using the Cytoscape (3.10.0). The EnrichmentMap plug-in on Cytoscape was used to visualize the GSEA KEGG (FDR≤0.01) and Reactome (FDR≤0.05) pathways for building the network graphs. Additionally, pathway analysis was performed using the Metascape for differentially upregulated genes in a test group compared to the reference group ^(25)^ (min expr 3, FDR≤0.05).

### RNA isolation and qPCR

RNA was extracted from a homogenized stomach biopsy in TRIzol (ThermoFisher Scientific) and precipitated from the aqueous phase using an equal volume of 70% ethanol. The mixture was transferred to an RNA isolation column (Omega Bio-Tek), and the remaining steps were followed according to the manufacturer’s recommendations. Quantitative RT-PCR was performed using the iTaq Universal Probes master mix (BioRad), and PCR was run on a QuantStudio 4 (ThermoFisher Scientific). The follow Taqman primer probes were used (ThermoFisher Scientific): *Nr3c1* (Mm00433832_m1)*, Tnf* (Mm00443258_m1)*, Il6* (Mm00446190_m1)*, Ifng* Mm01168134_m1)*, Atp4b* (Mm00437657_m1)*, Bhlha15* (Mm00487695_m1)*, Tff2* (Mm00447491_m1)*, Cftr* (Mm00445197_m1)*, Wfdc2* (Mm00509434_m1). *Ppib* (Mm00478295_m1) was utilized as a reference gene.

### Western Blot

Peritoneal macrophages were isolated as described above. Adherent cells were lysed in sodium dodecyl sulfate (SDS) sample lysis buffer (BioRad). Proteins were separated by gel electrophoresis on a 10% Tris-Glycine gel (BioRad), transferred to nitrocellulose membrane, and stained with anti-GR antibodies (D8H2, Cell Signaling Technologies) and anti-Beta-actin antibodies (MAB1501, Millipore Sigma) overnight at 4oC. Following stringency washes, the blots were probed with fluorescent secondary antibodies and were imaged with an iBright 1500 (Thermo Fisher Scientific).

### Statistical Analysis

GraphPad Prism 10 was used for the statistical analysis. An unpaired student T-test was used to compare two groups. One-way ANOVA with Tukey, a multiple comparisons test, was used when comparing more than two groups. Two-way ANOVA with Sidak’s multiple comparisons test was used when testing more than one variable (Figure 1C). A P-value of ≤0.05 was considered statically significant. All error bars are standard deviation ± the mean.

**Figure 1.**
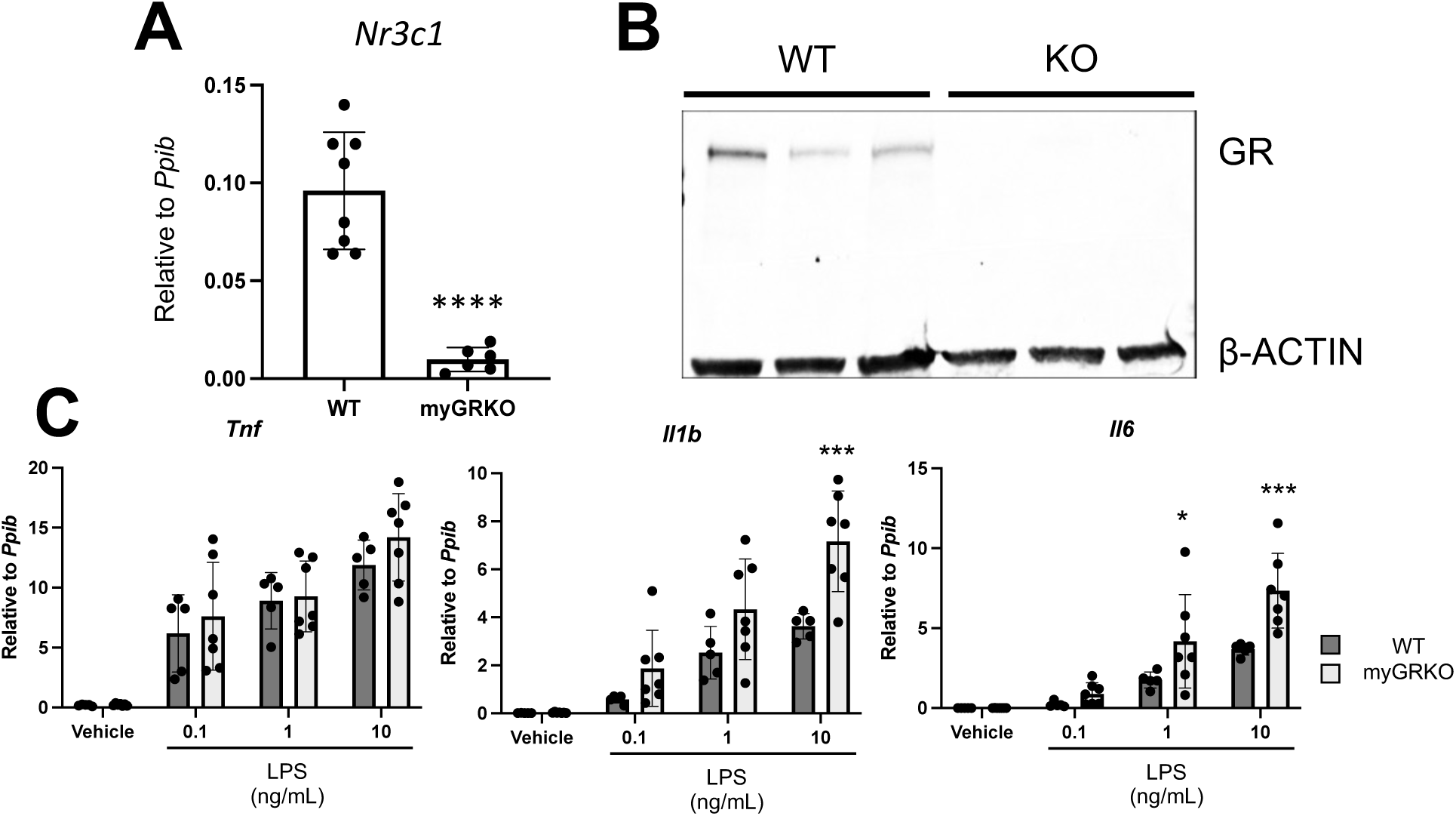
Loss of the glucocorticoid receptor poises macrophages to respond more aggressively to proinflammatory stimuli. (A) Quantitative RT-PCR for *Nr3c1,* (B) and Western blot for the glucocorticoid receptor (GR) using RNA (A) or protein (B) isolated from adherent cells 24 hours after isolation from Thioglycollate-induced peritoneal macrophages from myGRKO mice or WT littermate controls. (C) qRT-PCR of the indicated genes using RNA isolated from WT or GRKO peritoneal macrophages stimulated with the indicated amount of LPS for 6 hours. n≥6 (A) or ≥3 (B-C). \**P*≤0.05, \*\*\**P*≤0.001, and \*\*\*\**P*≤0.0001.

### Study Approvals

All mouse experiments were approved by the West Virginia University Animal Care and Use Committee.

## Results

### Glucocorticoid receptor deletion poises macrophages to respond more aggressively to proinflammatory stimuli

To investigate the impact of glucocorticoid signaling on the macrophage response to proinflammatory stimuli, we deleted the GR from the myeloid lineage by crossing GR floxed mice (exon 3) with the *Lysm* promoter-driven Cre recombinase ^(3)^. To confirm the deletion of the GR, Thioglycollate-induced peritoneal macrophages were isolated from myGRKO mice and WT littermate controls. Quantitative RT-PCR revealed that *Gr* transcript levels were significantly reduced in KO macrophages compared to WT controls (Figure 1A). Similarly, western blot revealed abundant GR protein in WT macrophages, but protein was not detectable in KO macrophages (Figure 1B). We examined how glucocorticoid signaling during monocyte-to-macrophage differentiation affected their subsequent activation. Thioglycollate-induced peritoneal macrophages were isolated from WT and myGRKO mice and cultured for 24 hours in steroid-free conditions, followed by a 6-hour stimulation with increasing concentrations of lipopolysaccharide (LPS). Relative transcript levels of the proinflammatory cytokines *Tnf, Il1b,* and *Il6* were measured by qRT-PCR. At a low LPS dose (0.1ng/mL), there were no significant differences between the cytokine response of WT and GRKO macrophages (Figure 1C). However, at moderate and high LPS doses, GRKO macrophages responded with significantly higher levels of *Il1b* and *Il6*, while *Tnf* levels were not significantly different at any LPS dose (Figure 1C). These findings suggest that signaling by endogenous glucocorticoids during macrophage differentiation has lasting effects on macrophage sensitivity to proinflammatory stimuli and that GRKO macrophages are poised toward a hyper-cytokine response to higher doses of proinflammatory stimuli.

### Unstimulated GRKO macrophages have increased chromatin accessibility in regions adjacent to proinflammatory genes

Our result demonstrates that GRKO macrophages have increased transcription of proinflammatory cytokines when compared to WT macrophages cultured in steroid-free media. Glucocorticoid signaling through the GR is best known for its transcriptional activities. However, recent evidence indicates that the GR signaling also shapes chromatin accessibility ^(29)^. Next, we tested whether the loss of glucocorticoid signaling altered chromatin accessibility. Thioglycollate-induced myGRKO and WT littermates received an intraperitoneal injection of 10^7^ *H. pylori* (Figure 2A). Three hours after treatment, macrophages were isolated from the peritoneal lavage by flow cytometry, and the chromatin landscape and transcriptome were assessed by ATACseq and RNAseq, respectively. Disruption of glucocorticoid signaling significantly affected the macrophage chromatin landscape from vehicle-treated mice, with 317 increased and 220 decreased differentially annotated regions (DARs) in vehicle-treated KO macrophages compared to vehicle-treated WT controls. Homer motif analysis of the increased DARs revealed increased accessibility of AP1 family transcription factors such as JUNB and FOS (Figure 2B). Genes that were spatially associated with DARs were annotated by the Genomic Regions Enrichment of Annotations Tool (GREAT) and assessed by Gene Ontology pathway analysis, revealing significant activation of inflammation-associated pathways such as inflammatory bowel disease, T cell activation, and chemokine production (Figure 2C). Next, RNAseq analysis revealed subtle differences in the transcriptomes of vehicle-treated WT and GRKO macrophages with 180 differentially expressed genes (DEGs) using a 0.05 *P-*value cutoff and only 27 DEGs when using a 0.01 *P-*value cutoff. Gene set enrichment analysis (GSEA) of expressed genes demonstrated enrichment of several proinflammatory pathways in GRKO macrophages using the Reactome database. Cytoscape summary of the GSEA results demonstrated that “Cytokine singling” was the most enriched node in GRKO macrophages (Figure 2D). These data indicate that glucocorticoid signaling significantly impacts the macrophage chromatin landscape, with increased accessibility in regions adjacent to proinflammatory genes. Moreover, these results suggest that resting GRKO macrophages are poised toward a hyperactive response.

**Figure 2:**
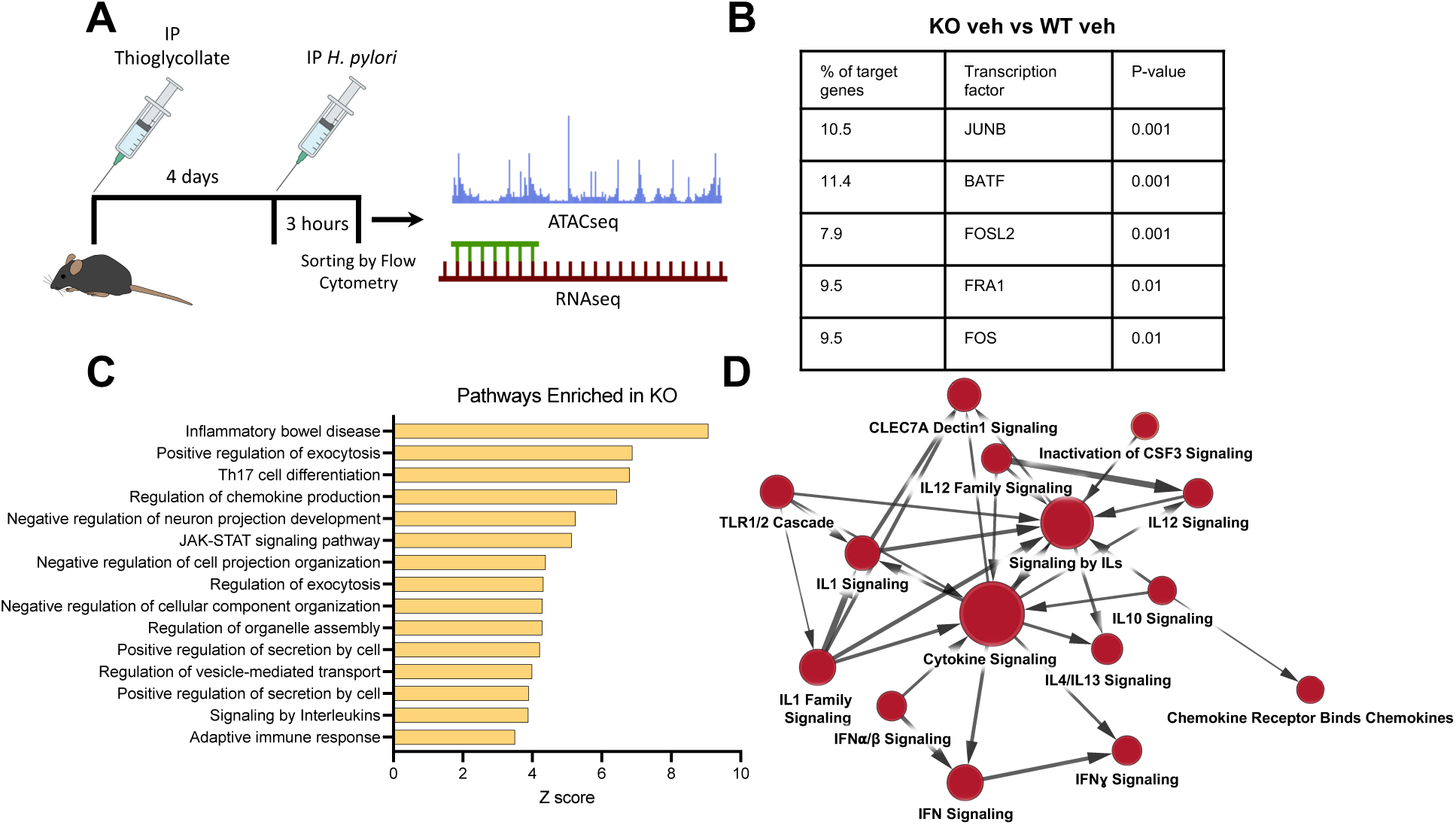
GRKO macrophages have increased chromatin accessibility in regions adjacent to proinflammatory genes. (A) Schematics showing the peritoneal challenge with *H. pylori* and subsequent macrophage sorting (B-C) Genomic regions that were significantly (p≤0.05) more accessible in vehicle treated GRKO macrophages were annotated by GREAT. (B) Homer motif analysis identified TF consensus sequences within these increased accessible regions. (C) Pathway analysis of the GREAT-annotated genes ranked by enrichment score. (D) Cytoscape Enrichment Map summarizing the Gene Set Enrichment Analysis of RNAseq of vehicle-treated WT and GRKO macrophages. Red dots indicate enrichment in GRKO macrophages.

### Glucocorticoid signaling in macrophages is required for their normal activation to bacterial infection

Signaling by endogenous glucocorticoids is critical for normal macrophage function. Previous studies have found that GR deletion from the myeloid compartment leads to increased mortality to LPS sepsis challenge ^(30)^, while others have reported impaired cytokine production and wound healing in models of Myocardial infarction ^(31)^. Our results indicate that in vivo GRKO macrophages may be poised to respond more aggressively to an inflammatory stimulus. Next, we examined how disruption of glucocorticoid signaling affected the macrophage response to an acute *H. pylori* challenge. As described above, peritoneal macrophages were isolated from WT and GRKO mice 3 hours after IP injection of *H. pylori* (Figure 2A). ATACseq revealed that *H. pylori* treatment induced massive chromatin reorganization, exacerbating the baseline differences between vehicle-treated macrophages with 3204 increased and 1260 decreased DARs in KO macrophages compared to WT *H. pylori*-treated macrophage controls. Homer motif analysis of the increased DARs indicated a massive increase in binding sites for AP1-associated transcription factors (Figure 3A). Gene ontology analysis of GREAT-annotated genes was used to describe the DARs between stimulated WT and KO macrophages. Within the top 15 pathways were annotations such as fever generation, prostaglandin synthesis, and regulation of macrophage activation (Figure 3B). These enriched pathways suggested that the loss of glucocorticoid signaling exacerbates proinflammatory gene expression. Perplexingly, though, 4 of the top 5 pathways were related to increased TGFβ signaling, suggesting parallel activation of anti-inflammatory gene networks.

**Figure 3:**
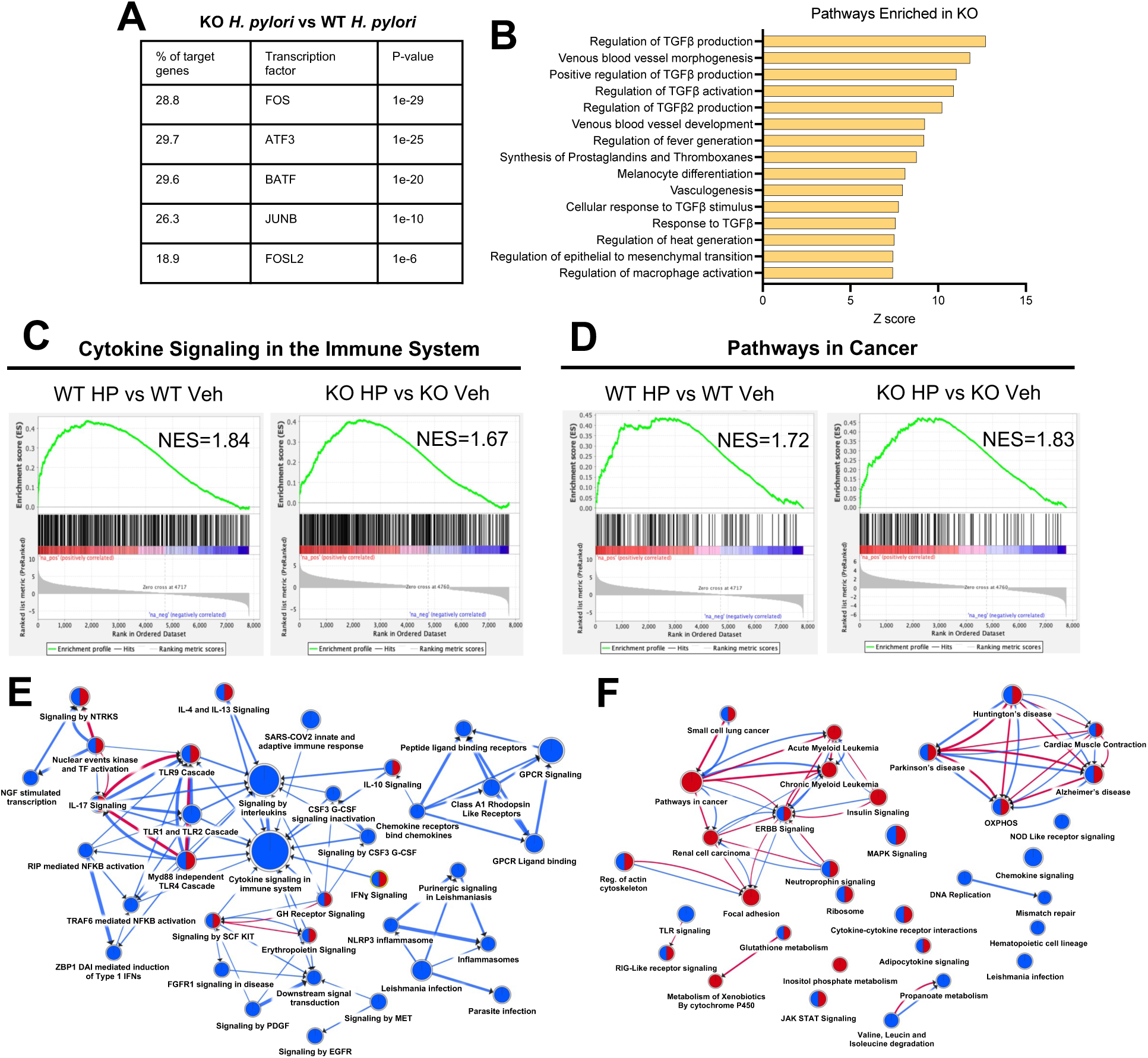
Glucocorticoid signaling in macrophages is required for their normal activation to inflammatory stimuli. (A-B) Genomic regions that were significantly (q≤0.01) more accessible in *H. pylori* challenged GRKO macrophages were annotated by GREAT. (A) Homer motif analysis identified TF consensus sequences within these increased accessible regions. (B) Pathway analysis of the GREAT-annotated genes ranked by enrichment score. (C) Gene Set Enrichment Analysis (GSEA) of the indicated Reactome (C) and KEGG (D) pathway comparing enrichment in activated WT and GRKO macrophages. (E-F) Cytoscape Enrichment Map summarizing the KEGG (E) and Reactome (F) GSEA results. Blue dots indicate enrichment in WT and red dots indicate enrichment in GRKO macrophages. Mixed dots indicate enrichment in both groups.

Next, we compared how *H. pylori* challenge affects the macrophage transcriptomes. *H. pylori* challenge caused significant enrichment of inflammatory pathways in both WT and GRKO macrophages when compared to their respective vehicle-treated controls. However, GSEA revealed that inflammatory-associated pathways such as “cytokine signaling in the immune system” were more enriched in WT macrophages compared to KO (normalized enrichment score, NES, 1.84 vs. 1.67) (Figure 3C). Cytoscape analysis of the enriched pathways from the Reactome and Kyoto Encyclopedia of Genes and Genomes (KEGG) databases indicated that activation of inflammatory gene sets was impaired in the *H. pylori-*challenged GRKO macrophages (Figure 3D-E). Interestingly, several KEGG cancer-associated gene sets were significantly enriched in GRKO macrophages (Figures 3D and 3F). These data demonstrate that the disruption of glucocorticoid signaling in macrophages causes an altered chromatin landscape and impaired transcription of inflammatory mediators. These data suggest that glucocorticoid signaling is required for normal macrophage activation and that the loss of glucocorticoid signaling leads to cancer-associated transcriptional responses and may lead to macrophage dysfunction.

### Loss of glucocorticoid signaling in the myeloid compartment blunts the gastric inflammatory response to H. pylori infection

Our transcriptomics analysis indicated that the loss of glucocorticoid signaling impaired macrophage activation. Therefore, we next examined how myGRKO mice responded to gastric *H. pylori* colonization. WT and myGRKO mice were infected with *H. pylori,* and inflammation within the gastric corpus was assessed 2 months post-colonization. Flow cytometry was used to determine gastric immune infiltration (Figure 4A). Tissue-resident leukocytes were similar in vehicle-treated WT and myGRKO stomachs (Figure 4B). *H. pylori* colonization induced significant gastric leukocyte infiltration, but infiltration was significantly reduced in myGRKO stomachs (Figure 4B). Within the myeloid compartment, macrophage recruitment, but not neutrophils, was significantly blunted in myGRKO mice compared to WT controls (Figure 4C). Interestingly, T cell recruitment was also significantly impaired in myGRKO mice, suggesting that normal macrophage function is critical for coordinating the T cell response to *H. pylori* infection. Quantitative RT-PCR was used to measure *Tnf, Il6,* and *Ifng* mRNA levels within the gastric corpus mucosa. All three proinflammatory cytokines were significantly increased in WT *H. pylori-*infected mice (Figure 4D). While all three cytokines also increased in myGRKO stomachs, *Il6* and *Ifng* did not achieve statistical significance. These data demonstrate that glucocorticoid signaling in the myeloid compartment is critical for mounting an immune response to *H. pylori* infection and provides increasing evidence that GRKO macrophages are dysfunctional.

**Figure 4:**
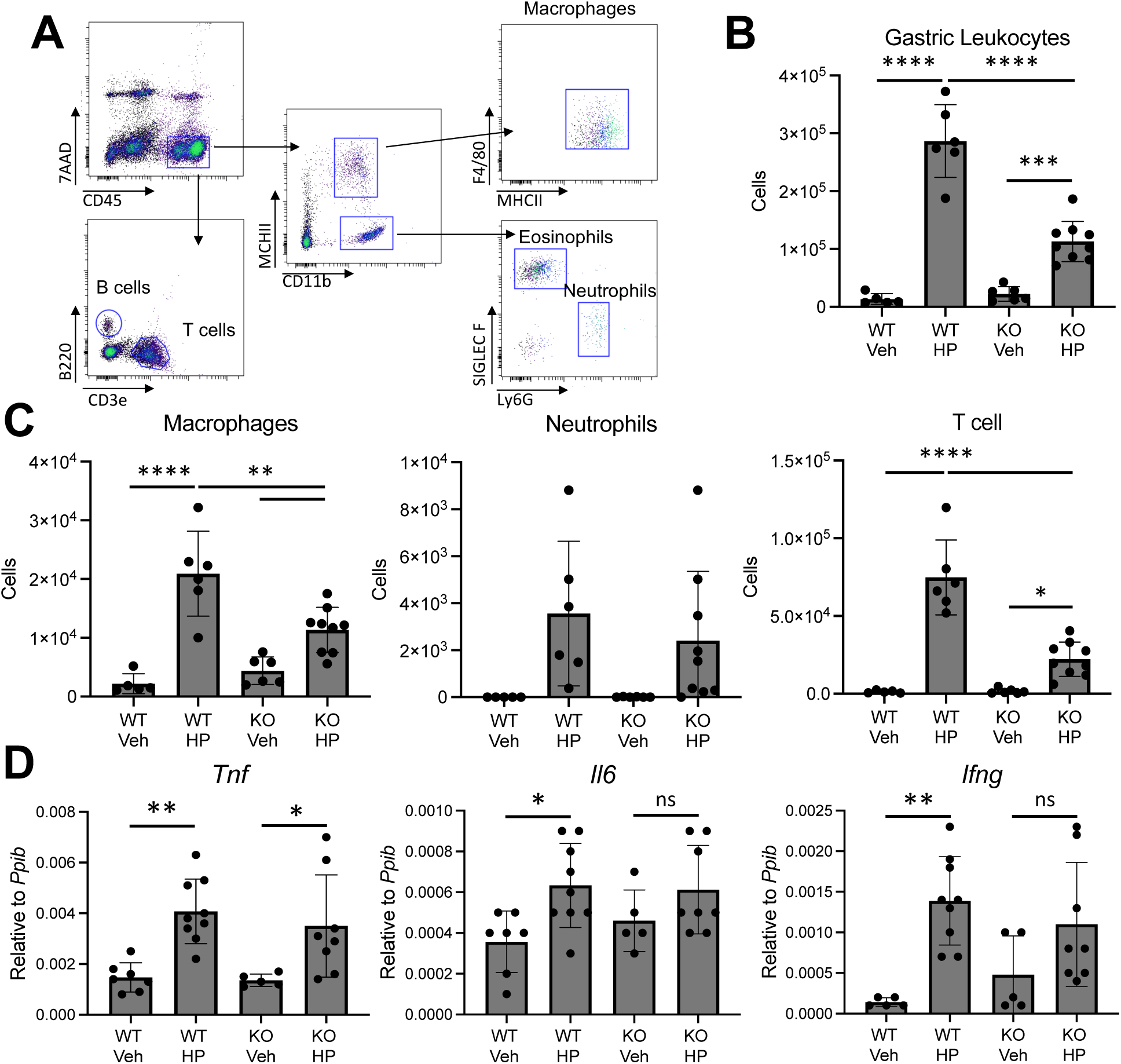
Loss of glucocorticoid signaling in the myeloid compartment impairs the gastric inflammatory response to *H. pylori* infection. (A) Representative flow cytometry plots demonstrating the gating strategy. (B-C) Flow cytometry analysis of the indicated cell types within the gastric corpus 2 months after vehicle treatment or *H. pylori* infection. (D) Quantitative RT-PCR of the indicated genes within the gastric corpus 2 months after vehicle treatment or *H. pylori* infection. n≥5. \**P*≤0.05, **P≤0.01, \*\*\**P*≤0.001, and \*\*\*\**P*≤0.0001.

### myGRKO mice are protected from Helicobacter-induced gastric atrophy and metaplasia

Chronic inflammation in response to *H. pylori* infection damages the gastric epithelium, driving metaplasia development and gastric cancer initiation. We found that the loss of glucocorticoid signaling in macrophages impairs their activation and blunts gastric T-cell recruitment (Figure 4). Next, we investigated how glucocorticoid signaling affected *the development of Helicobacter-*induced gastric atrophy and metaplasia. The *H. pylori* PMSS1 strain is poorly immunogenic ^(32)^ and induces limited gastric atrophy 2 months post-infection (Figure 5A-B). In WT mice, *H. pylori* infection induced mild atrophy of the gastric corpus glands, but gastric atrophy was largely absent in the gastric corpus of *H. pylori-*infected myGRKO mice. Quantitative RT-PCR of the lineage markers *Atp4b* (parietal cells), *Mist1* (mature chief cells), and *Tff2* (mucous neck cells) demonstrated that *H. pylori-*induced gastric atrophy was extremely limited 2 months post-colonization (Figure 5B).

**Figure 5:**
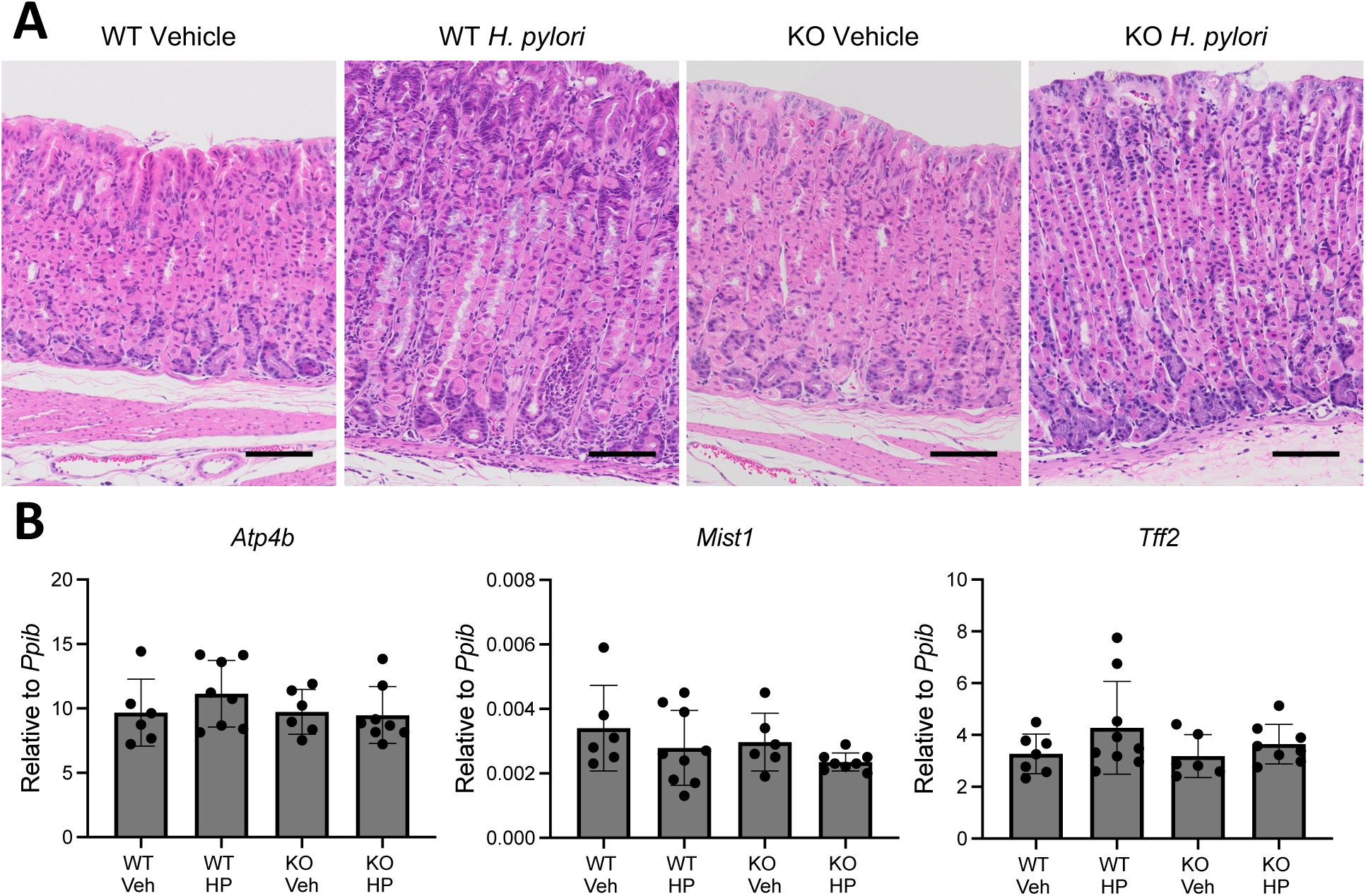
myGRKO mice are protected from *Helicobacter*-induced gastric atrophy. (A) Representative H&E micrographs of the gastric corpus greater curvature. Scale bars=100µM. (B) Quantitative RT-PCR of parietal cell (left), chief cell (center), and mucous neck cell (right) lineage markers utilizing RNA from the gastric epithelial corpus. n≥6.

We next infected mice with the natural mouse pathogen *Helicobacter felis*, which is well-known to induce severe gastric inflammation, atrophy, and metaplasia^(33, 34)^. *H. felis* infection induced significant inflammation and gastric corpus remodeling in WT and myGRKO mice (Figure 6A). However, gastric atrophy was more extensive in WT mice, with a complete absence of chief cells, a significant reduction of parietal cells, and a massive expansion of GSII+ mucous cells (Figure 6A-B). Moreover, qRT-PCR demonstrated significant loss of chief and parietal cell transcripts and expansion of mucous neck cell transcripts (Figure 6C). In contrast, gastric atrophy was notably more subdued in *H felis-*infected myGRKO mice (Figure 6A-B). Parietal cell loss and GSII+ mucous cell expansion were reduced compared to WT-infected controls (Figure 6A-B). Parietal and mucous neck cell transcript levels were not significantly different from myGRKO vehicle controls. Finally, we examine how the infection affected pyloric metaplasia development. In WT mice, *H. felis* infection caused widespread expression of CD44v9, extending to the base of gastric corpus glands (Figure 7A). Moreover, there was a significant increase in the PM transcripts *Cftr* and *Wfdc2* (Figure 7B). In contrast, CD44v9 expression was modestly reduced in myGRKO stomachs, and *Cftr* and *Wfdc2* expression was reduced (Figure 7A-B). These results demonstrate that macrophage dysfunction impairs the gastric inflammatory response to *Helicobacter* infection and indicates that glucocorticoid signaling in macrophages is required for their normal responsiveness to bacterial infection.

**Figure 6:**
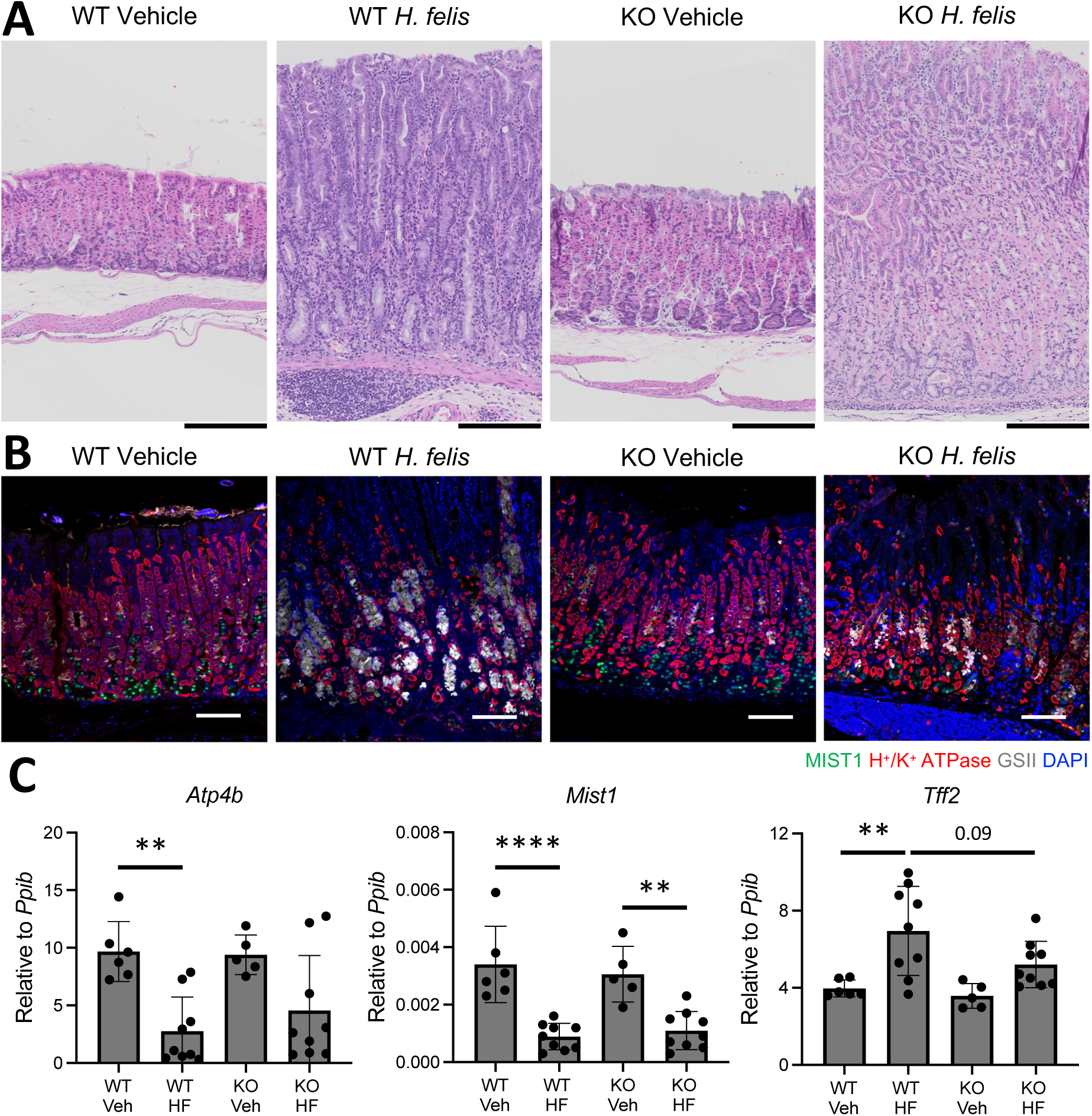
myGRKO mice are protected from the *H. felis* induced atrophic gastritis. Tissues and RNA were isolated from the gastric corpus greater curvature of WT and myGRKO 6 months post-*H. felis* colonization. (A) Representative H&E micrographs. Scale bars=200µM (B) Representative immunostaining. Sections were probed with antibodies against the MIST1 (mature chief cells, green) the H+/K+ ATPase (parietal cells, red), and the *Griffonia simplicifolia* lectin (mucous neck cells, grey). Nuclei were stained with DAPI. Scale bars=100µM. (**C**) Quantitative RT-PCR of RNA from the gastric corpus for the indicated genes. n≥5. **p≤0.01; ****p≤0.0001.

**Figure 7.**
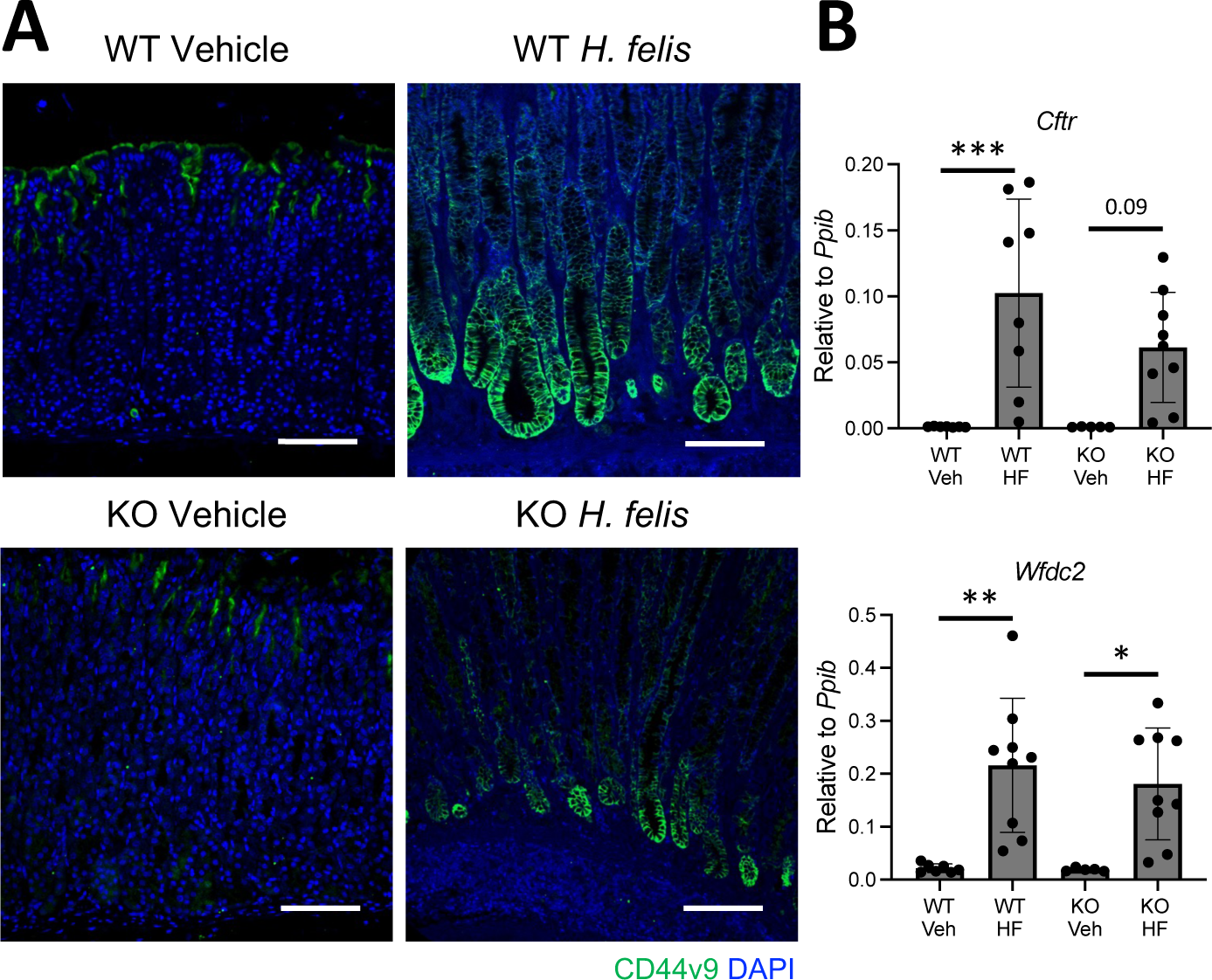
Pyloric metaplasia is blunted in *H. felis-*infected myGRKO stomachs. Tissues and RNA were isolated from the gastric corpus greater curvature of WT and myGRKO 6 months post-*H. felis* colonization. (A) Representative immunostaining for the pyloric metaplasia marker CD44v9. Nuclei are stained with DAPI. Scale bars=100µM. (B) Quantitative RT-PCR of the indicated pyloric metaplasia marker genes. n≥5. *p≤0.05; **p≤0.01, ***p≤0.001.

## Discussion

Our data demonstrate that glucocorticoid signaling is required for the normal macrophage response to *H. pylori*. Loss of glucocorticoid signaling caused reorganization of the chromatin landscape, increasing accessibility in regions adjacent to proinflammatory genes, both in resting and stimulated macrophages. Upon challenge with *H. pylori,* GRKO macrophage activation was impaired, and their transcriptional response was skewed towards cancer-associated gene sets. Within the stomach, myGRKO mice exhibited defective immunity against *H. pylori* infection, with fewer inflammatory cells and reduced expression of proinflammatory cytokines. Finally, when myGRKO mice were infected with the more immunogenic bacteria *H. felis*, they exhibited lower levels of gastric atrophy. Together, these results suggest that glucocorticoid signaling poises macrophages to respond to pathogens and demonstrates that glucocorticoid signaling is required for normal macrophage activation. Moreover, we found that defective macrophage activation impairs the overall gastric inflammatory response to *H. pylori* infection.

Glucocorticoids have been clinically used for over half a century and remain one of the most widely used drugs for combating inflammatory diseases ^(35)^. There is a formidable body of literature on the anti-inflammatory effects of glucocorticoid treatments. Their wide-ranging immunomodulatory effects are achieved through cell-type-specific transcriptional changes ^(36)^. The effects of glucocorticoid treatment are primarily anti-inflammatory, where they oppose proinflammatory cytokine transcription, inhibit B cell receptor and T cell receptor signaling, suppress pattern recognition receptor expression and activation, oppose antigen presentation, etc ^(1)^. However, many of the immunomodulatory effects of glucocorticoids are dictated by ligand concentration, and endogenous glucocorticoids play a more nuanced role in regulating immune activation. In macrophages, high doses of glucocorticoids strongly suppress the transcription of proinflammatory genes, while low doses promote the transcription of proinflammatory genes ^(4)^. Similarly, low doses of glucocorticoids delivered before an inflammatory challenge enhanced the macrophage response to LPS ^(37)^, suggesting that glucocorticoids are important for preparing the immune system for noxious stimuli.

In this study, we avoided the complication of dosing animal subjects by exposing macrophages to inflammatory stimuli in vivo. Thioglycollate-induced macrophages are primarily monocyte-derived ^(38)^, and WT cells were differentiated in vivo under the influence of endogenous glucocorticoid, whereas GRKO macrophages were insensitive to the direct effects of endogenous corticosteroids. In a previous study, Bhattacharyya et al. reported that myGRKO mice died from septic shock when challenged with LPS, while WT controls survived the challenge ^(30)^. Thus, our initial expectation was that GRKO macrophages would develop a hyperactive response to an intra-peritoneal *H. pylori* challenge. However, GRKO macrophages exhibited defective activation of proinflammatory genes and instead exhibited enrichment of several cancer-associated gene networks. Moreover, myGRKO mice mounted a weaker gastric inflammatory response to *H. pylori* colonization. These results demonstrate a dichotomy in the immunomodulatory effects of endogenous glucocorticoids. Bhattacharyya et al. reported that endogenous glucocorticoids are critical for limiting inflammatory intensity to a high dose of LPS and preventing septic shock ^(30)^. In contrast, our results indicate that, during steady-state conditions, endogenous glucocorticoids function to prime the macrophage response and are critical for effective immunity.

Glucocorticoid signaling impacts a wide array of cellular responses, and it is estimated that glucocorticoids regulate up to 20% of expressed genes ^(39)^. However, the cell-type-specific transcriptional changes regulated by glucocorticoids are primarily determined through pre-existing chromatin accessibility patterns ^(40)^. Ligand-activated GR recruits several co-activators and co-repressors that affect the chromatin landscape, including histone methyltransferases and acetyltransferases as well as the SWI/SNF chromatin remodeling complex ^(29, 41)^. We found that *H. pylori* stimulation induced dramatic differences in the chromatin landscape in WT vs GRKO macrophages. Interestingly, many of the increased DARs in KO macrophages contained AP1 binding motifs, a transcription factor complex that drives the expression of many proinflammatory cytokines. The AP1 complex is a well-known target of GR trans-repression, and the increased accessibility of AP1 motifs is likely a direct effect of the loss of the GR ^(42)^. Interestingly, pathway analysis of the increased DARs in GRKO macrophages indicated a striking activation of TGFβ signaling. Similar findings were reported by Galuppo et al. who found that *Tgfb1* expression was increased in GRKO macrophages ^(31)^. TGFβ is an anti-inflammatory cytokine that suppresses macrophage activation and promotes alternative macrophage polarization ^(43)^. The increase in TGFβ signaling may represent a compensatory mechanism to oppose the hyperactivation caused by the loss of glucocorticoid signaling.

Within the stomach, macrophages act as a double-edged sword. During steady-state conditions, they are the most abundant tissue-resident leukocytes in the gastric corpus, where they maintain tissue homeostasis ^(17)^. However, monocyte-derived macrophages produce damaging cytokines following gastric injury that exacerbate tissue and promote pyloric metaplasia development ^(14, 15)^. During *H. pylori* infection, macrophages simultaneously control bacterial loads and contribute to gastric atrophy. Activation of proinflammatory M1-like macrophages is associated with reduced bacterial colonization but also increased epithelial damage and gastric atrophy ^(44, 45)^. In contrast, factors that impair macrophage activation lead to increased bacterial loads and reduced gastric atrophy ^(11, 46)^. Our results demonstrate that impaired macrophage activation through deletion of the GR reduced gastric atrophy and PM development.

The direct effects of macrophages and macrophage-derived cytokines on gastric atrophy are unclear. We have previously shown that systemic glucocorticoid depletion induces spontaneous gastric inflammation and PM and that macrophage depletion, either through clodronate liposomes or blocking monocyte infiltration in *Cx3cr1* KO mice, prevents these gastric pathologies ^(15)^. Similarly, work by Peterson et al. showed that macrophage-derived cytokines drive PM development following L635-induced gastric injury ^(13, 14)^. While these studies demonstrate that macrophages contribute to gastric atrophy progression, others have shown that Helicobacter-induced gastric atrophy is blocked in T cell-deficient mice ^(47)^. Thus, macrophage effects on *Helicobacter-*induced gastric atrophy may occur through coordinating the T cell responses. Supporting this notion, macrophage depletion in *H. pylori-*infected gerbils reduced lymphocytes infiltration and tertiary lymphoid tissue development ^(48)^. Here, we found that myGRKO mice had reduced gastric T cells. Thus, in addition to the cell-intrinsic macrophage dysfunction caused by defective glucocorticoid signaling, these effects also impair adaptive immunity.

In summary, glucocorticoid signaling is critical for multiple phases of the immune response, preparing the immune system to combat pathogens effectively while limiting the immune response of pathogenically activated immune cells. Our results show that endogenous glucocorticoid signaling impacts the macrophage transcriptome and chromatin landscape and that the loss of glucocorticoid signaling induces macrophage dysfunction. While this study focused on cells with either intact or disrupted glucocorticoid signaling, the temporal effects of glucocorticoid signaling have important clinical relevance and were not addressed here. In addition, our findings underscore the critical role that gastric macrophages play in the immune response to *H. pylori,* where they impact the T cell response and promote gastric atrophy and metaplasia development. These findings, combined with previous reports ^(15, 17, 49)^, demonstrate that glucocorticoid signaling is important for regulating gastric immunity and may impact gastric cancer risk.

## Abbreviations

WT: Wild Type
GR: Glucocorticoid receptor
GRKO: Glucocorticoid Receptor Knock Out
PM: Pyloric metaplasia
lipopolysaccharide: LPS
PAMPs: Pathogen-associated molecular patterns
PMSS1: Pre-mouse Sydney Strain 1
GS II: *Griffonia simplicifolia II*
FACS: Fluorescence activated cell sorting
ATAC: Assay for Transposase-accessible Chromatin
DAR: Differentially Annotated Regions
GREAT: Genomics region enrichment of annotations
GSEA: Gene Set Enrichment Analysis
KEGG: Kyoto Encyclopedia of Genes and Genomes
qRT-PCR: quantitative real time polymerase chain reaction

## Author Contributions

S.K. and J.T.B planned the study. S.K., S.A.D. and J.T.B performed experiments. X.X., L.W., and G.H. analyzed genomics data. S.K. and J.T.B analyzed all data and drafted the manuscript. All authors reviewed and approved the final manuscript.

